# Biased sampling confounds machine learning prediction of antimicrobial resistance

**DOI:** 10.1101/2025.01.07.631773

**Authors:** Yanying Yu, Nicole E Wheeler, Lars Barquist

## Abstract

Antimicrobial resistance (AMR) poses a growing threat to human health. Increasingly, genome sequencing is being applied for the surveillance of bacterial pathogens, producing a wealth of data to train machine learning (ML) applications to predict AMR and identify resistance determinants. However, bacterial populations are highly structured and sampling is biased towards human disease isolates, meaning samples and derived features are not independent. This is rarely considered in applications of ML to AMR. Here, we demonstrate the confounding effects of sample structure by analyzing over 24,000 whole genome sequences and AMR phenotypes from five diverse pathogens, using pathological training data where resistance is confounded with phylogeny. We show resulting ML models perform poorly, and increasing the training sample size fails to rescue performance. A comprehensive analysis of 6,740 models identifies species- and drug-specific effects on model accuracy. We provide concrete recommendations for evaluating future ML approaches to AMR.

## Introduction

Antimicrobial resistance (AMR) is a severe threat. Currently, an estimated 4.95 million deaths are associated with AMR each year^1^, and this number is only predicted to increase. Urgent action is required to prevent further escalation, slow epidemic spread, and preserve our capacity to administer advanced medical therapies relying on surgery or immunosuppression. Surveillance based on whole genome sequencing is playing an increasingly important role in monitoring the emergence and spread of AMR^2^. Taking full advantage of this accumulating data will require automated systems that can transform genome sequences into actionable intelligence, namely predictions of resistance profiles and identification of new mechanisms of resistance^3–5^. Machine learning (ML) approaches have the potential to fill this niche, though a number of challenges remain in implementing such systems^6^.

One underappreciated challenge is in the assumptions made by ML methods themselves. Most classical ML methods assume training data are independent and identically distributed, which is not true of pathogen surveillance samples due to the underlying structure of bacterial populations. During epidemic spread, successful clones spread rapidly. If this spread is due in part to acquisition of AMR determinants it could lead to an association between the phenotype and phylogenetic markers that do not directly contribute to AMR. These non-causative associations are likely further exacerbated by biased sampling focused on human disease in high-income countries^5,7^ leaving large regions of the phylogeny unexplored. In the context of microbial genome-wide association studies (GWAS), these phylogenetic effects have been mitigated in part by the application of mixed-effect models that attempt to correct for sampling biases^8–10^. To date, the implications of biased sampling for ML approaches to AMR prediction have not been systematically investigated, despite the clear risk of confounding^11^. Here, by constructing realistic pathological scenarios, we show that biased sampling combined with population structure can dramatically affect the performance of ML methods for predicting AMR. We investigate the factors underlying the performance of ML methods, and provide concrete recommendations for the future evaluation of ML approaches to predicting AMR.

### Sampling effects bias machine learning prediction of AMR

To comprehensively evaluate the impact of population structure in predicting AMR, we collected between 3,204 and 7,188 genomes for three gram-negative species (*Escherichia coli, Klebsiella pneumoniae, Salmonella enterica*) and two gram-positive species (*Staphylococcus aureus, Streptococcus pneumoniae*) representative of current WHO priority pathogens^12^. In total, our dataset includes resistance phenotypes for 27 antibiotics each represented by over 1,000 resistant genomes (**FIG S1**). As a representative ML model we used LightGBM^13^, a gradient boosted decision tree method representative of the tree ensemble-based methods that generally perform well for predicting AMR^14,15^. We defined discrete clades on the phylogenetic tree for each species (**FIG 1A&B, FIG S2-6A**) based on deep divergences between branches.

Based on the clade structure of our data, we developed pathological training cases simulating biased sampling. In our first approach, Scheme A (**FIG 1C**), we selected two clades, and a balanced test set was held out from one (the test clade). We then devised 4 scenarios, representing biased sampling. Scenarios a and b excluded either resistant or sensitive isolates from training data for the test clade, respectively. As a positive control, we also included scenarios c and d, which excluded sensitive or resistant strains from the non-test clade. Compared to a random split of training and test data, models trained on biased datasets exhibited much lower area under the receiver operating characteristic curve (AUC) scores (**FIG 1D**). Exclusion of resistant strains led to low precision and high recall of sensitive strains (**FIG S7A**), while the opposite was observed for exclusion of sensitive strains (**FIG S7A**), indicating that the ML models were conflating indicators of lineage with genuine indicators of AMR. This was not observed in our control scenarios, and, importantly, persisted even when both resistant and sensitive strains were present in the test clade, but at biased proportions (**FIG S7BC**).

**FIG 1.**
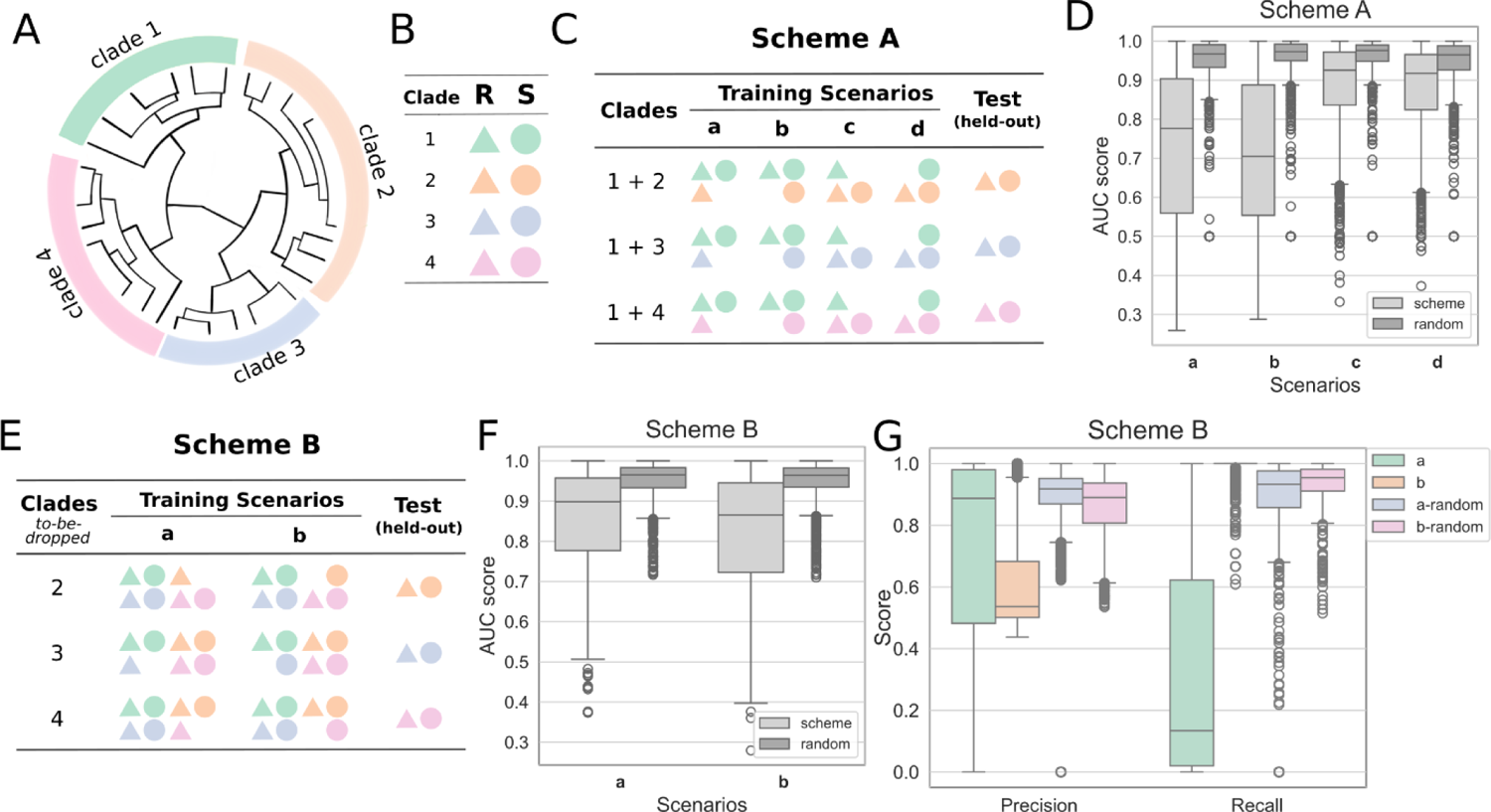
Population structure hinders the surveillance of emerging AMR strains. (A) An illustration of defining clades for genomes in one species. (B) Legend for panels C and E, R: resistant (triangles), S: susceptible (circles). (C) An example of training scenarios for scheme A based on the clade splitting in A. Models were trained on clade 1 and one of the other clades, such as clades 1 and 2, clades 1 and 3, and clades 1 and 4. Four training scenarios were tested for each antibiotic and species. In each scenario, either resistant or susceptible samples from one of the paired clades were excluded from the training data, while the held-out genomes from the clade that is not clade 1 were used as the test sets. (D) The area under the ROC curve (AUC) scores in each scenario for scheme A for all antibiotics and species. Light grey: train-test split followed scheme A, dark grey: random train-test split. (E) An example of training scenarios for scheme B based on the clade splitting in A. Models were trained on all samples except for resistant or susceptible samples from one clade. The held-out genomes from the same clade were used as the test sets. (F) The AUC scores for each scenario for scheme B for all antibiotics and species. Light grey: train-test split followed scheme B, dark grey: random train-test split. (G) The precision and recall scores for scheme B; a-random and b-random indicate testing on random train-test splits of the pathological unbalanced data.

It is often implicitly assumed that simply increasing sample sizes is sufficient to improve ML models. To test this, we developed a second approach, Scheme B, where we additionally included training data from remaining clades, while still excluding resistant or sensitive isolates from the test clade (**FIG 1E**). This did lead to an improvement in the AUC compared to Scheme A (**FIG 1F**). However, extreme biases remained in both precision and recall (**FIG 1G**), indicating that trained models continued to conflate lineage and AMR markers even in the presence of a larger training data set.

### AMR predictivity varies across antibiotics and species

To further understand the factors driving model performance, we trained a random forest meta-model to predict the AUC scores of all trained models (**Table S2**), including species, training scenario, clade-wise distance, sample size, and drug class as predictors. We then interpreted this meta-model using Shapley additive explanation (SHAP) values^16^ (**FIG 2A**).

**FIG 2.**
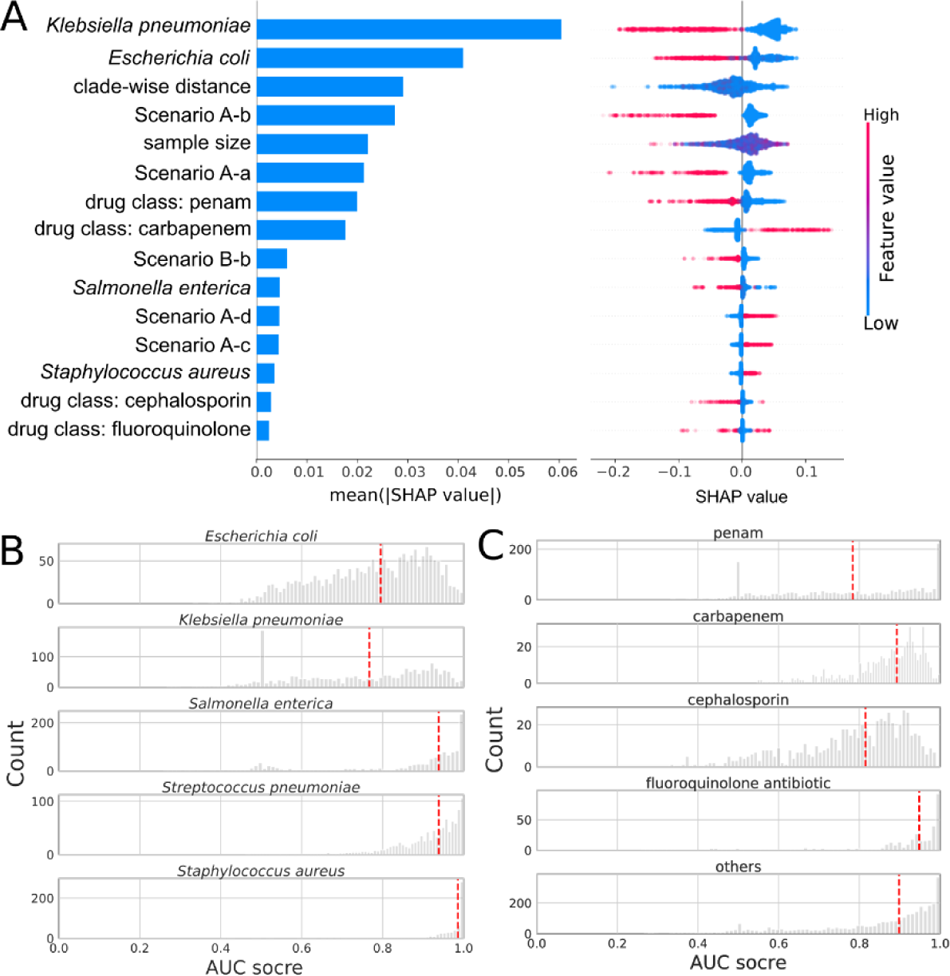
AMR predictivity varies across antibiotics and species. (A) SHAP values for the top 15 features from a random forest model trained on AUC scores from both schemes A and B for all five species. Global feature importance is given by the mean absolute SHAP value (left), while the beeswarm plot (right) illustrates feature importance for each guide prediction. (B) The distribution of AUC scores for each species. Red dashed lines indicate the median values. (C) The distribution of AUC scores for four drug classes shown in A. Red dashed lines indicate the median values.

This analysis reaffirmed the inferior performance associated with our pathological training scenarios. Additionally, this analysis showed that training on data sourced from Gram-negative species yielded diminished AUC scores (**FIG 2B**), presumably due to the presence of the outer membrane^17^, and that drug class also had large and at times contradictory effects on model performance (**FIG 2C**). Given the impact of the species on model performance, we asked how the model interpretation differs when training on each species (see **FIG S2-6C**). As an example, the effects of fluoroquinolone varied in three species (**FIG S2C, S4C, S5C**) and also in the model encompassing all species (**FIG 2A**). Subsequent interpretation of models trained on each clade for ciprofloxacin (**FIG S9**) further underscored the distinct determinants for prediction across clades and species. Among the known genes associated with ciprofloxacin resistance^18^, the gene *parC* prominently featured in most clades of *E. coli* and *S. aureus.* In contrast, the gene *gyrA* was exclusive to a single clade within *S. aureus.* In other clades of *E. coli* and *S. enterica*, the strongest predictors diverged, encompassing genes such as transcriptional regulator *kdgR*, mobilization gene *mobB*, quinone oxidoreductase *qorA*, and genes encoding hypothetical proteins.

To test if this lack of common features was general, we trained clade-specific models for all drug classes. Surprisingly, minimal feature overlap was identified, both within the top 10 features and when summing the top features based on SHAP values to account for 50% of total SHAP values (**FIG S8A&B**). This scarcity of common predictive genes or SNPs across clades may in part be a result of the small number of strong predictors (**FIG S8C**), and provides a partial explanation for the strong effects we observed for our pathological training scenarios.

### Implications for AMR prediction

ML-based modeling of AMR can serve multiple purposes. These include at least two major applications: 1) prediction of strain resistance, and 2) identification of causal variants. While these two applications are related, achieving one does not necessarily imply achieving the other. If biased sampling reflects biological reality, i.e., all strains of a particular lineage truly are resistant to an antibiotic, then models that conflate lineage markers with resistance may still provide accurate predictions for surveillance purposes. This can be seen for instance in applications that have used phylogenetic placement to predict AMR^19^. However, such models will not provide insights into the underlying molecular mechanisms of resistance, and will fail on strains from previously unseen lineages or resistant strains emerging from a former sensitive lineage. Conversely, identifying causal variants requires disentangling lineage effects from true resistance determinants, a task complicated by the structure and biases present in genomic datasets.

Our results demonstrate that current ML approaches are particularly vulnerable to the confounding effects of biased sampling and population structure. Even when sample sizes are increased, models struggle to generalize beyond the specific clades included in the training set, underscoring the limitations of scaling as a solution to bias. This is supported by previous work that has shown phylogenetic structure is an important feature for predicting AMR^14^, and that many AMR prediction tools perform poorly when tested on phylogenetically-constructed benchmark sets^20^. The lack of overlap we found in predictive features across clades further highlights the challenges in identifying universal resistance markers, with most predictions being dependent on the phylogenetic background. These findings are consistent with previous observations in bacterial GWAS, where lineage effects are a persistent source of confounding and require explicit correction.

Future efforts to improve ML-based AMR prediction must incorporate strategies to address these challenges. One potential approach is the development of lineage-aware ML algorithms that explicitly model the hierarchical structure of bacterial populations, borrowing ideas from mixed-effect models commonly used in bacterial GWAS^8–10^. Another promising direction is the integration of population-genomic frameworks that incorporate evolutionary signals, such as measures of phylogenetic conservation, into feature selection and model interpretation^21^. Additionally, benchmark datasets that include well-curated, globally representative samples are essential for evaluating and comparing new approaches, ensuring models are tested in realistic scenarios that capture the diversity encountered in clinical and environmental surveillance.

Importantly, sampling strategies must also evolve to support these developments. Expanding surveillance efforts to underrepresented regions and settings, particularly in low- and middle-income countries (LMICs), will provide more balanced datasets and reduce biases toward high-income countries and outbreak scenarios^7^. Targeted sampling that prioritizes diversity—both within and across clades—will be critical for developing models capable of generalizing across phylogenetic backgrounds. Beyond sampling, careful experimental design, including the use of balanced test sets and rigorous evaluation metrics such as those employed here, will further ensure that models are robust to confounding effects. Including algorithms based on simple assumptions, such as phylogenetic placement^19^, as baseline models for ML development should help in determining if machine learning is providing additional insights or simply recapitulating phylogenetic structure.

In conclusion, addressing the interplay between population structure and AMR prediction will require a multifaceted approach that includes improved sampling, algorithmic innovation, and systematic evaluation of proposed prediction methods. Only by confronting these challenges can we unlock the full potential of machine learning to provide actionable insights into AMR, advancing both our surveillance capabilities and our understanding of resistance mechanisms. These efforts will be critical to combating the global threat of AMR and ensuring the continued efficacy of life-saving antimicrobial therapies.

## Supplementary material

**FIG S1.**
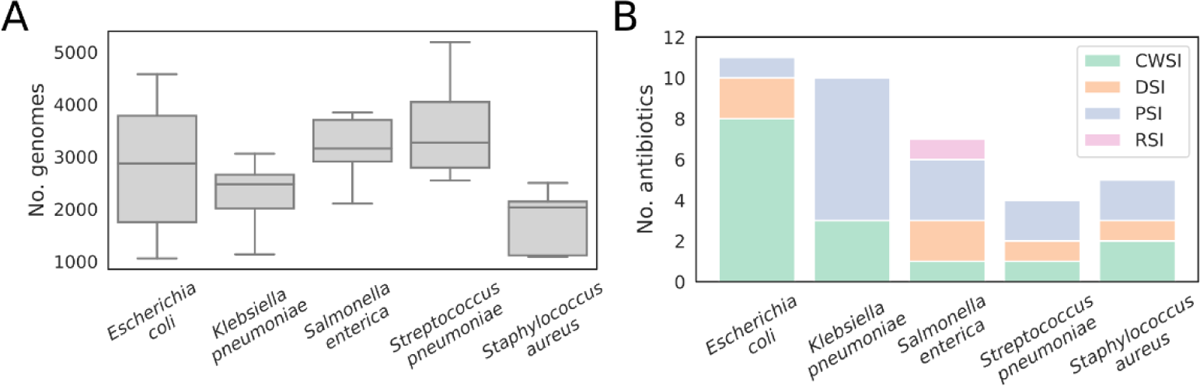
Overview of the collected data. (**A**) The number of genomes included in the training for different antibiotics in five species. (B) The number of antibiotics in each MOA in five species. CWSI: cell wall synthesis inhibitor, DSI: DNA synthesis inhibitor, PSI: protein synthesis inhibitor, RSI: RNA synthesis inhibitor. The full list of tested antibiotics for each of the five species is included in **Table S1**.

**FIG S2.**
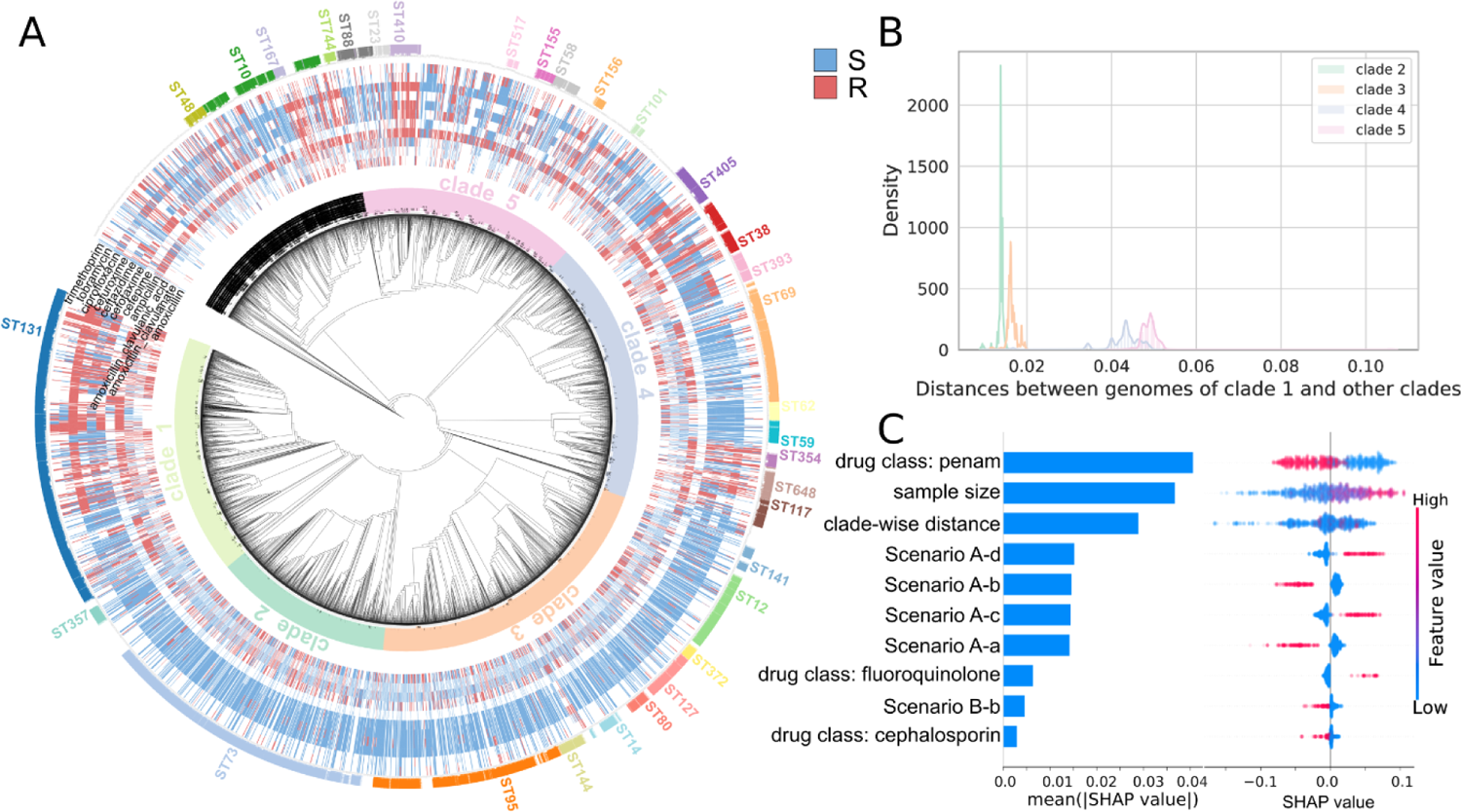
Phylogenetic tree and model interpretation in *Escherichia coli*. (**A**) Clade definition for model training, the antibiotic phenotypes, and the sequence types (ST) shown on the phylogenetic tree. S: susceptible, R: Resistant. (**B**) The distribution of pairwise distances between genomes of clade 1 and other clades. (**C**) SHAP values for the top 10 features from a random forest model trained on AUC scores from both schemes A and B for *Escherichia coli*.

**FIG S3.**
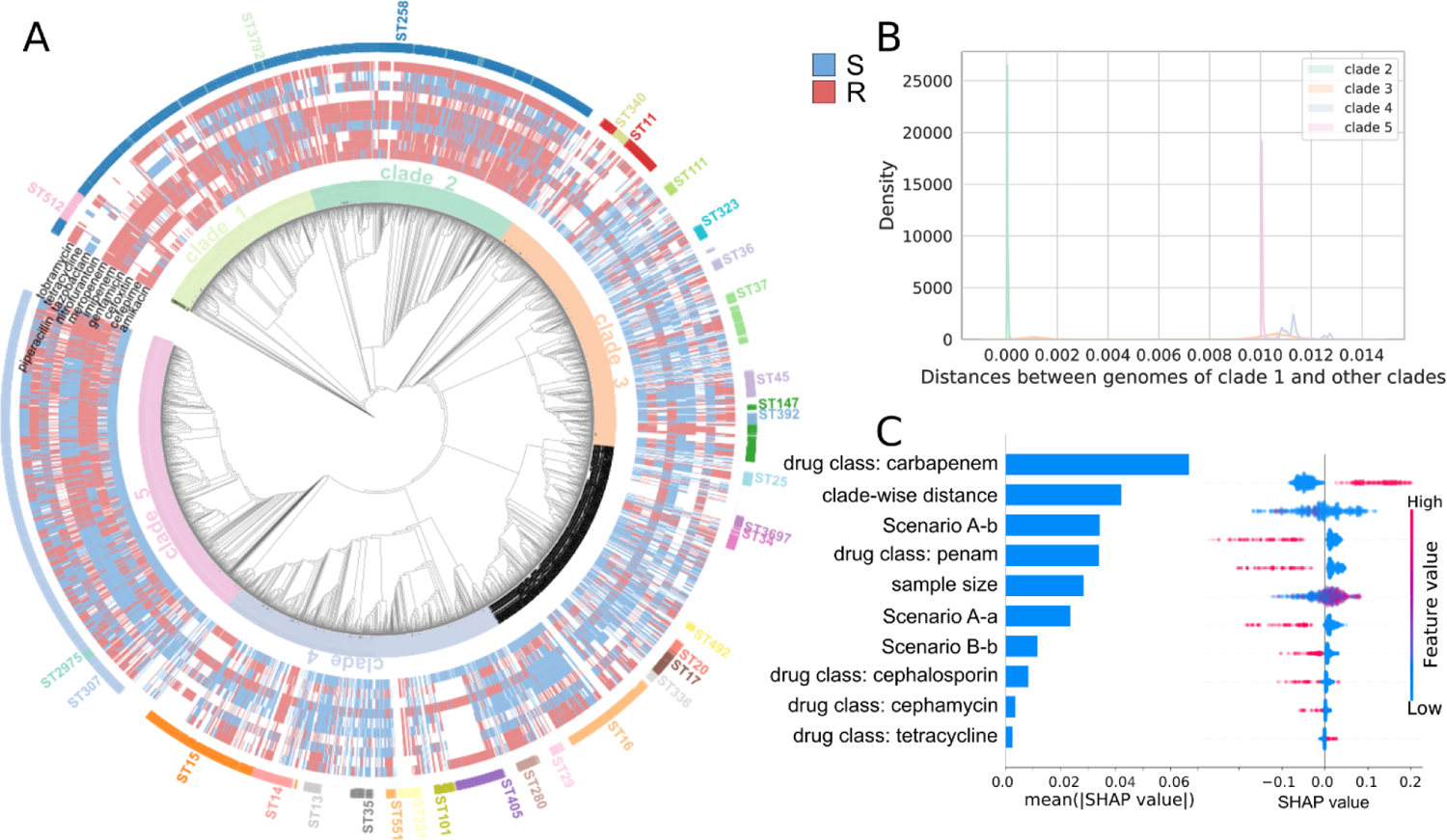
Phylogenetic tree and model interpretation in *Klebsiella pneumoniae*. (**A**) Clade definition for model training, the antibiotic phenotypes, and the sequence types (ST) shown on the phylogenetic tree. S: susceptible, R: Resistant. (**B**) The distribution of pairwise distances between genomes of clade 1 and other clades. (**C**) SHAP values for the top 10 features from a random forest model trained on AUC scores from both schemes A and B for *Klebsiella pneumoniae*.

**FIG S4.**
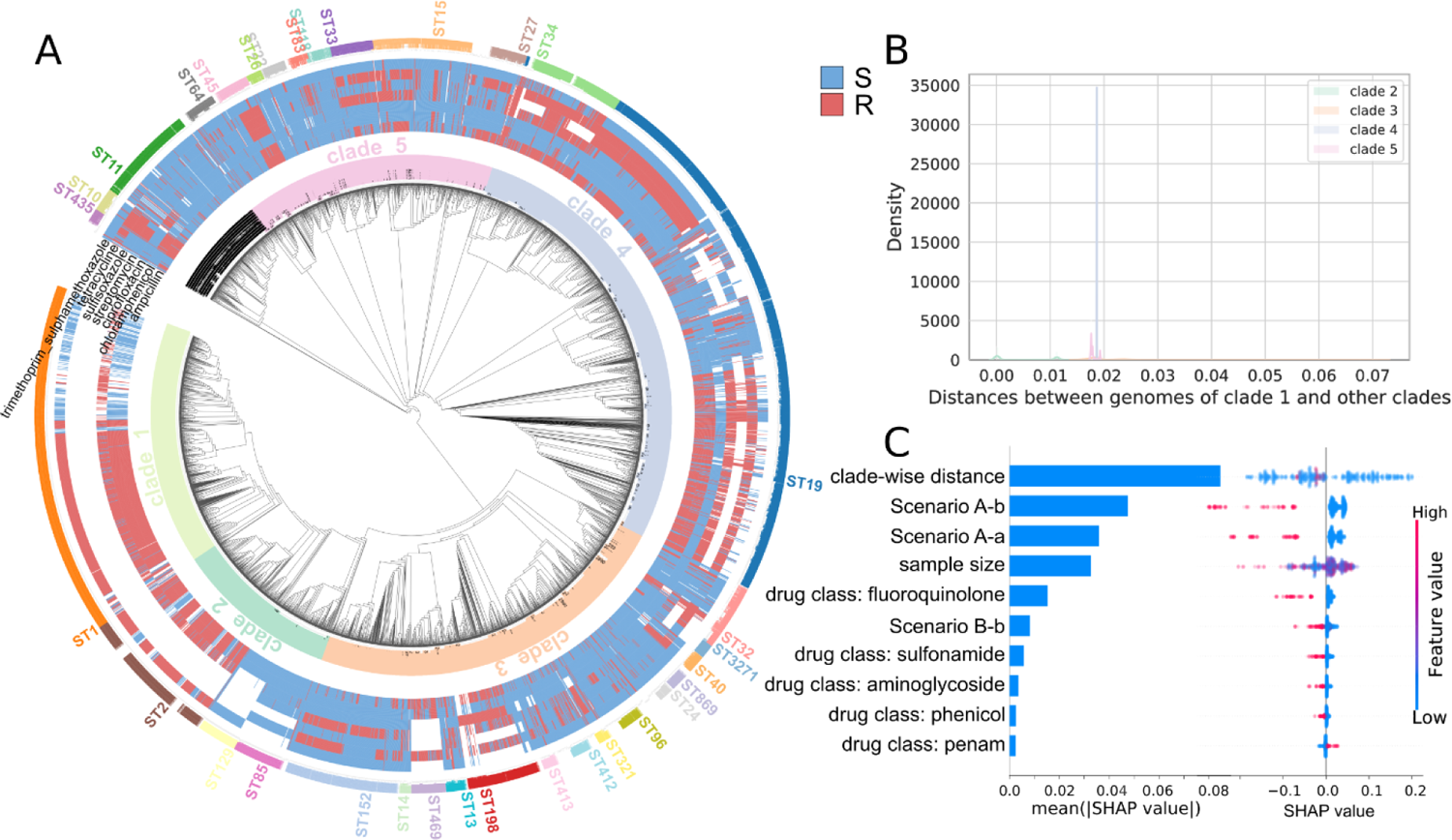
Phylogenetic tree and model interpretation in *Salmonella enterica*. (**A**) Clade definition for model training, the antibiotic phenotypes, and the sequence types (ST) shown on the phylogenetic tree. S: susceptible, R: Resistant. (**B**) The distribution of pairwise distances between genomes of clade 1 and other clades. (**C**) SHAP values for the top 10 features from a random forest model trained on AUC scores from both schemes A and B for *Salmonella enterica*.

**FIG S5.**
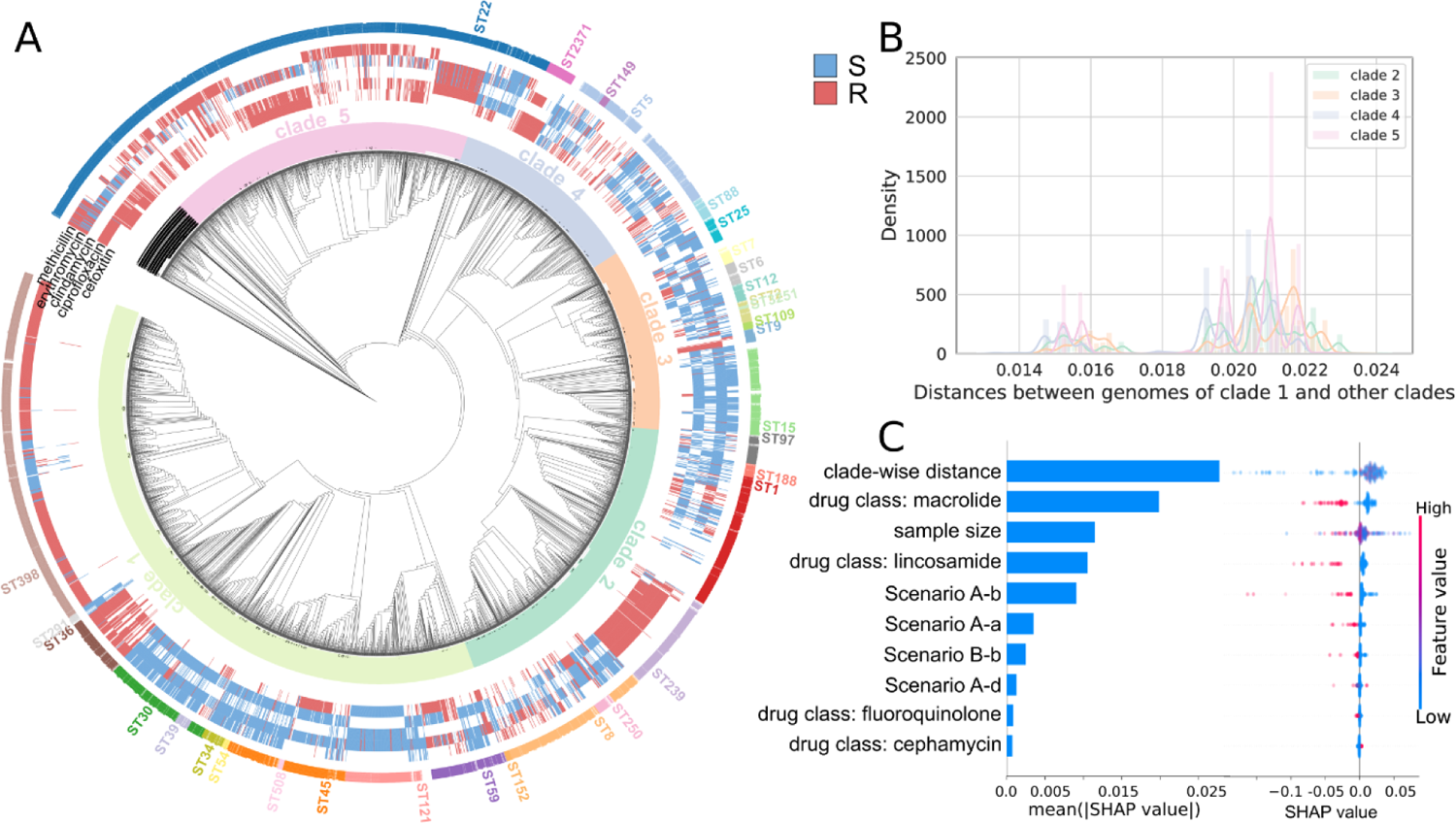
Phylogenetic tree and model interpretation in *Staphylococcus aureus*. (**A**) Clade definition for model training, the antibiotic phenotypes, and the sequence types (ST) shown on the phylogenetic tree. S: susceptible, R: Resistant. (**B**) The distribution of pairwise distances between genomes of clade 1 and other clades. (**C**) SHAP values for the top 10 features from a random forest model trained on AUC scores from both schemes A and B for *Staphylococcus aureus*.

**FIG S6.**
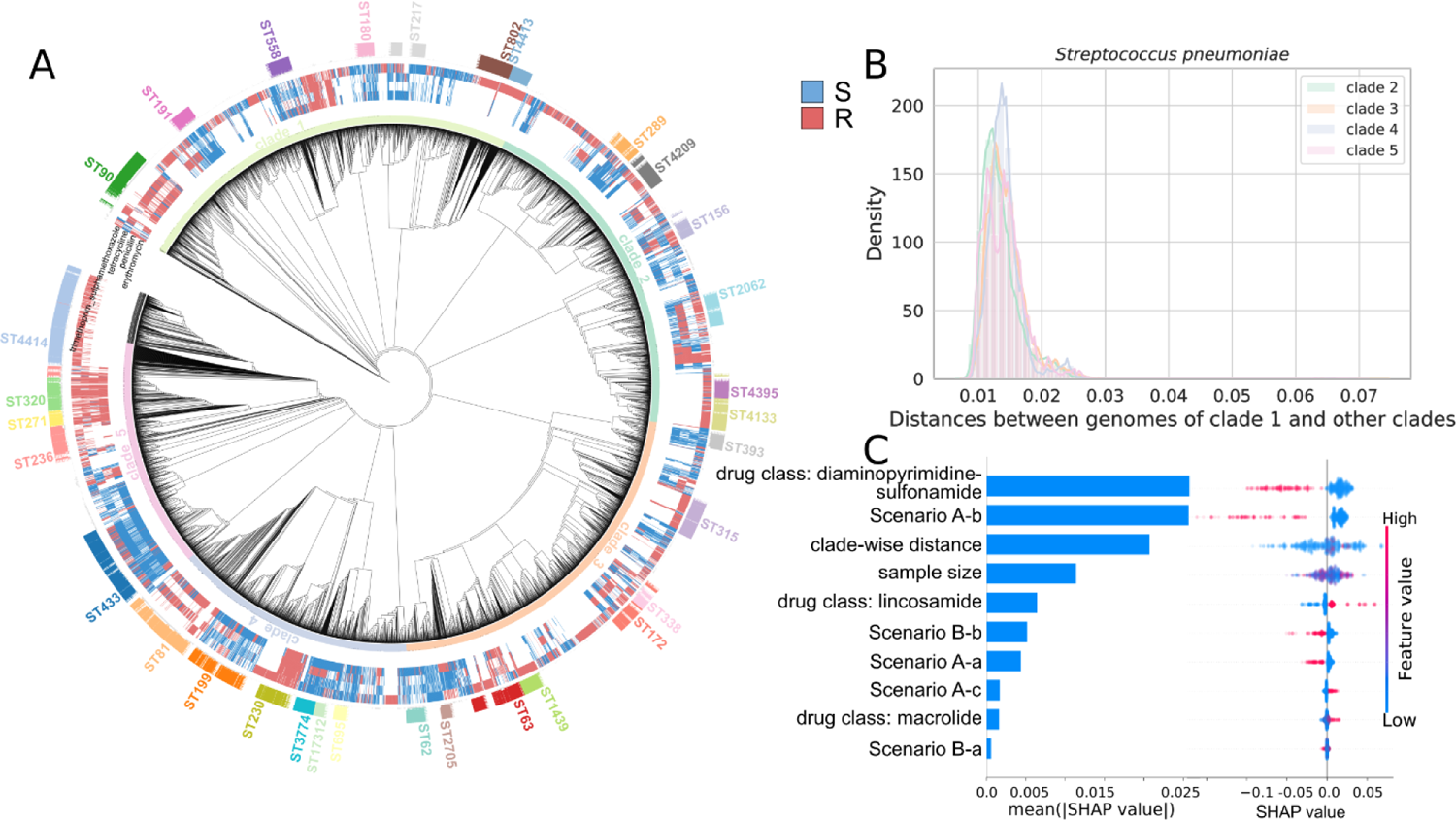
Phylogenetic tree and model interpretation in *Streptococcus pneumoniae*. (**A**) Clade definition for model training, the antibiotic phenotypes, and the sequence types (ST) shown on the phylogenetic tree. S: susceptible, R: Resistant. (**B**) The distribution of pairwise distances between genomes of clade 1 and other clades. (**C**) SHAP values for the top 10 features from a random forest model trained on AUC scores from both schemes A and B for *Streptococcus pneumoniae*.

**FIG S7.**
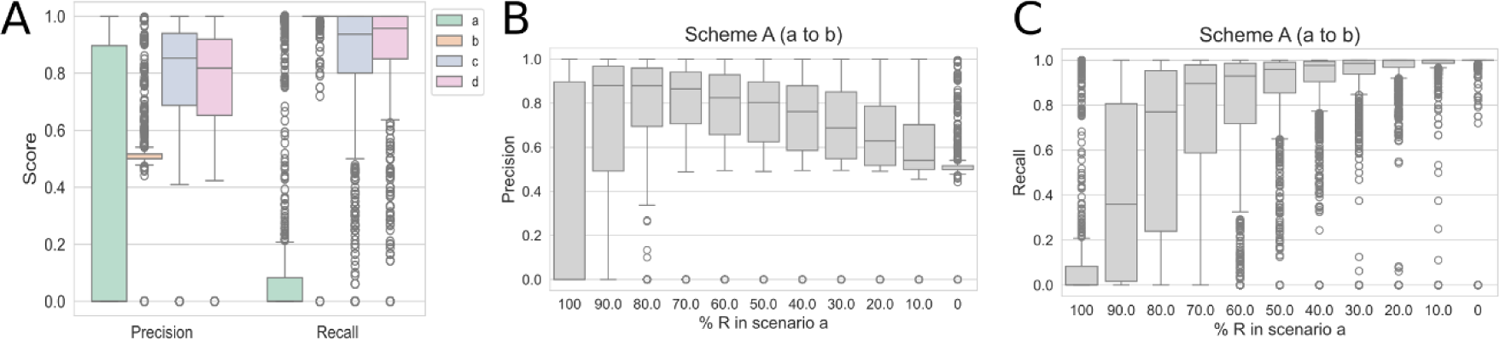
Precision and recall scores for scheme. **A.** (**A**) Scores in each scenario for all antibiotics and species. (**B**) Precision scores and (**C**) recall scores when decreasing resistant samples and increasing susceptible samples from the paired clade were included in training, with susceptible samples considered as positives.

**FIG S8.**
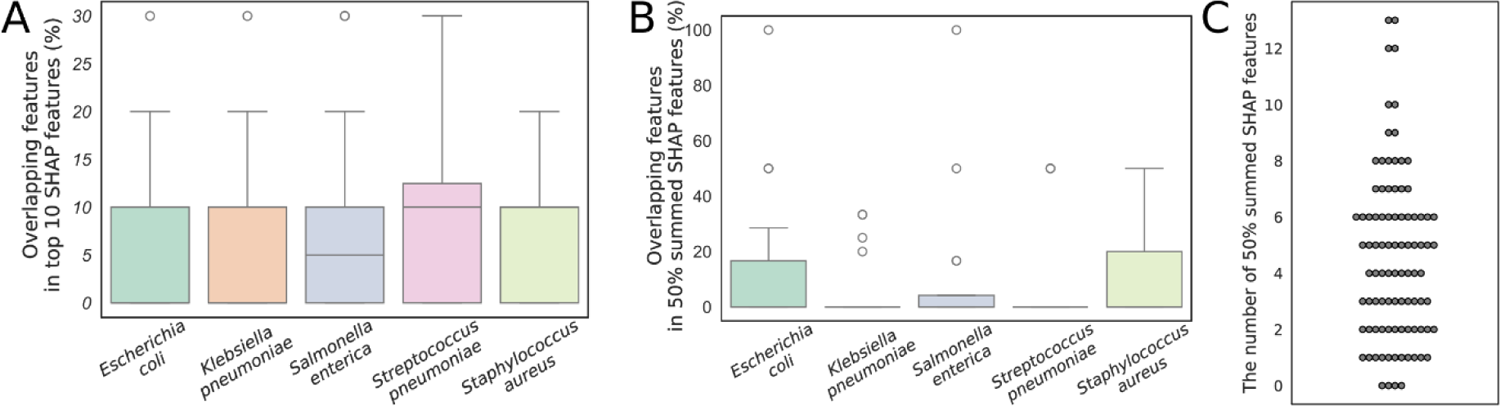
Overlapping predictive features across models trained on each clade in (**A**) top 10 features and (**B**) top features with SHAP values summed up to 50% of total SHAP values. (**C**) The distribution of the number of features with SHAP values that summed up to 50% of total SHAP values.

**FIG S9.**
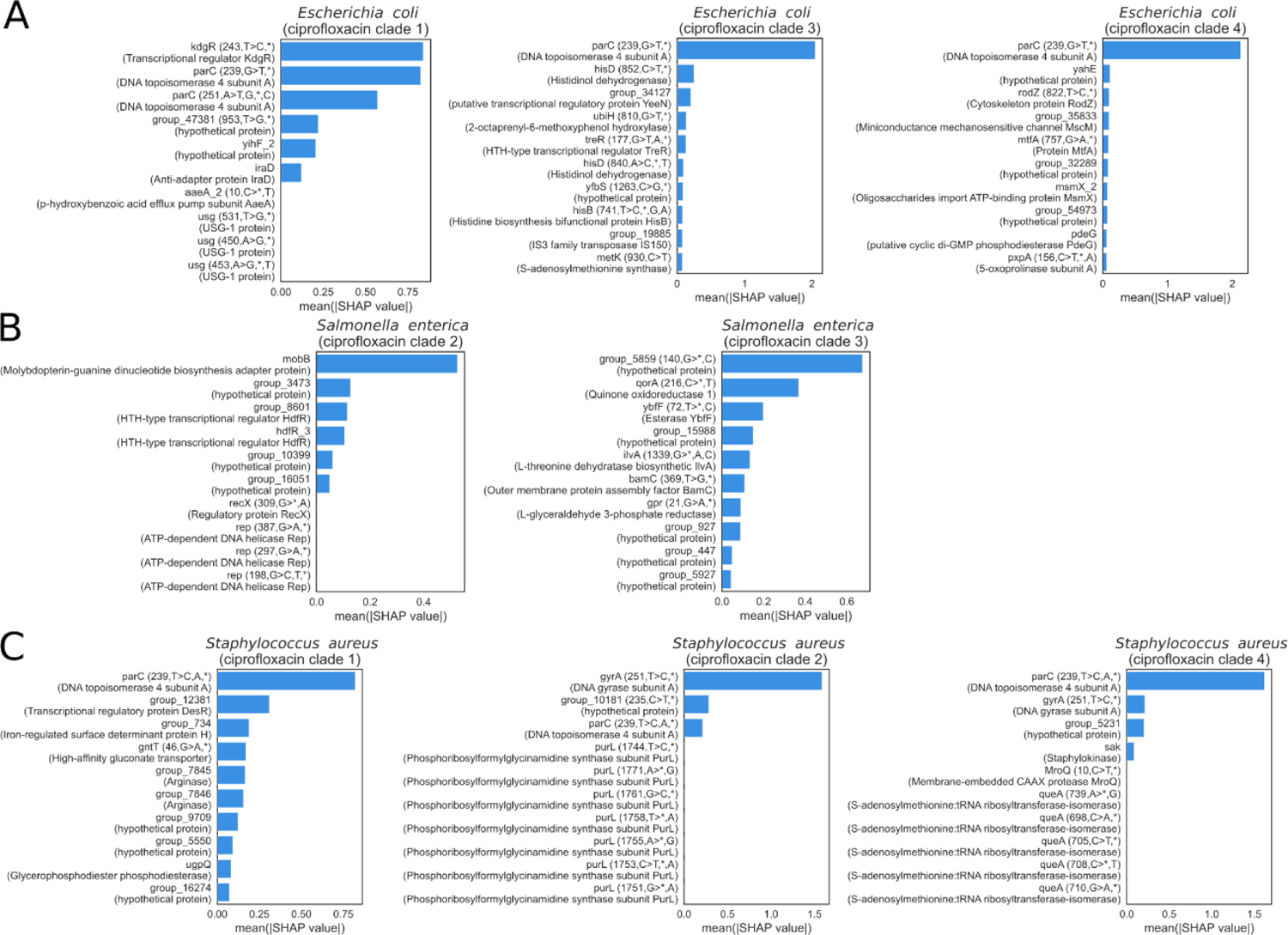
The interpretation of models trained on individual clades for ciprofloxacin in (**A**) *Escherichia coli*, (**B**) *Salmonella enterica,* and (**C**) *Staphylococcus aureus*. Clades with fewer than 50 resistant or susceptible strains were excluded.

## Methods

### Data collection

Whole genome sequences and antibiotic phenotypes were extracted from ^14^ and ^22^. 1,681 *Escherichia coli* genomes from ^14^ were assembled from FASTQ files using Velvet (Version 1.2.10) ^23^ with a hash length of 45. Genome quality was checked using CheckM (version 1.2.1)^24^ with the identical species name for the parameter “taxon_set species”. Genomes with Completeness lower than 95% or Contamination higher than 5% were removed from subsequent analysis.

### Phylogenetic analysis

Genomes were annotated using Prokka (version 1.14.6) ^25^. Annotation files were used as input for core genome alignment using Roary (version 3.7.0) ^26^ with a 95 minimum percentage identity for all genomes of each species. SNP-sites (version 2.5.1) ^27^ was used for SNP-calling on the core genome alignments. The phylogenetic tree was built on core genome alignments using IQ-TREE (version 2.2.0.3) ^28^ with the GTR model. Pairwise genome distance was calculated using the tree file from IQ-TREE and “phylogeny_distance.py” from Pyseer (version 1.3.10) ^10^ with the “--lmm” parameter. Sequence types were determined based on genes on https://pubmlst.org/data. For *Escherichia coli*, the #1 set of genes was used. For the visualization of the phylogenetic tree and related information, iTol (version 6.8) ^29^ was used. Only the top 25 sequence types with the highest number of genomes were shown in the iTol plots.

### Applying machine learning to predict antibiotic resistance

#### Training target and model type

LightGBM (version 3.3.2)^13^ binary classification models were trained to predict resistant or susceptible phenotypes for each antibiotic of each species. Antibiotics with less than 1,000 genomes or with either resistant or susceptible samples higher than 80% were not included in the training.

#### Training samples

Genomes in each species were manually split into several clades based on the phylogenetic trees (**FIG. S2-6**). For each antibiotic, clades with either resistant or susceptible samples less than 50 were excluded. Two schemes were tested. In scheme A (**FIG 1C**), one clade was selected to be paired with each of the other clades. If clade 1 contained sufficient resistant or susceptible samples, then it would be selected. Otherwise, other clades were tested for sufficient resistant or susceptible samples one by one until one clade was selected. Samples from the selected clade and one pairing clade were included for each training. 20 resistant and 20 susceptible samples from the pairing clade were held out for testing. Four scenarios were tested in scheme A. In each scenario, either all resistant or susceptible samples from one clade were excluded. When testing for including resistant and susceptible samples in training, between 10 and 90% of resistant samples and 90 and 10% of susceptible samples from one clade were included. In scheme B (**FIG 1E**), two scenarios were tested. In each scenario, either all resistant or susceptible samples from one clade were excluded, while both resistant and sensitive samples from all other clades were included. For all scenarios, 10-fold cross-validation was applied to resistant and susceptible samples in each clade and 90% of the samples were included in each fold according to the specific scenario in each scheme. In the random train-test split regime for both schemes A and B, training samples remained the same while the test set composition depended on the training exclusions. For example, if resistant samples from clade N were excluded from training, only sensitive samples from clade N were included in the test set. For training models on each clade, all samples in the clade were included.

#### Training features

SNPs from SNP-sites and the presence and absence table from Roary were used as features. In each training, features with a variance equal to zero were removed.

#### Hyperparameter tuning

Hyperparameters for the LightGBM model were optimized using hyperopt (version 0.2.7) ^30^. Search space included: ‘max_bin’ from range 50 to 500 with a step of 50, ‘bagging_fraction’ from range 0.01 to 1, ‘bagging_freq’ from range 0 to 10, ‘feature_fraction’ from range 0.01 to 1, ‘subsample_for_bin’ from range 30 to 0.8*sample size, ‘max_depth’ from range 1 to 16 with a step of 1, ‘learning_rate’ from range 0.01 to 1, ‘lambda_l2’ from range 0 to 100, ‘min_data_in_leaf’ from range 1 to 300, ‘min_gain_to_split’ from range 0 to 15, ‘num_leaves’ from range 2 to 100 with a step of 1. All samples were used for tuning. The mean of the balanced accuracy score in 10-fold cross-validation was used as the scorer. And the optimized set of hyperparameters was used in scenarios in the same scheme.

#### Evaluation

The held-out samples and the excluded samples were combined and an equal amount of resistant and susceptible samples were randomly drawn for calculation. The mean values of the scores over 100 repetitions were used.

#### Applying machine learning to interpret model performance

The default random forest regression model from scikit-learn (version 1.0.2) ^31^ was used to predict the AUC scores of the models. Species, scenarios, clade-wise distance, sample size, and drug classes were included as features. Drug classes were defined according to CARD ^32^. The clade-wise distance was calculated as the mean of all genome pair-wise distances between two clades. Species, scenarios, and drug classes were one-hot encoded. When training on scores from one species, the feature species was removed.

#### Model interpretation

SHAP values were calculated using the ‘shap_values’ function in TreeExplainer (version 0.39.0)^16^ with all samples. SHAP value plots were generated with the ‘summary_plot’ function in shap.

## Code and data availability

All code necessary to reproduce the results in the manuscript is available at https://github.com/BarquistLab/AMR_prediction. Files and results of machine learning have been deposited at Mendeley and are available through the following URL: https://data.mendeley.com/datasets/zs2mbjv7dn/1.

## Notes

https://data.mendeley.com/datasets/zs2mbjv7dn/1

### Competing Interest Statement

The authors have declared no competing interest.

